# Extending Finlay-Wilkinson regression with environmental covariates

**DOI:** 10.1101/2022.12.14.520390

**Authors:** Hans-Peter Piepho

## Abstract

Finlay-Wilkinson regression is one of the most popular methods for analysing genotype-environment interaction in series of plant breeding and variety trials. The method involves a regression on the environmental mean, computed as the average of all genotype means. The environmental mean is an index for the productivity of an environment. Productivity is driven by a wide array of environmental factors. Increasingly, it is becoming feasible to characterize environments explicitly using quantitative measurements of these factors. Hence, there is mounting interest to replace the environmental index with an explicit regression on such observable environmental covariates. This paper reviews the development of such methods. The focus is on parsimonious models that allow replacing the environmental index by regression on synthetic environmental covariates formed as linear combinations of a larger number of observable environmental covariates. Two new methods are proposed for obtaining such synthetic covariates, which may be integrated into genotype-specific regression models. The main advantage of such explicit modelling is that predictions can be made also for new environments where trials have not been conducted. A published dataset is employed to illustrate the proposed methods.

## 1 INTRODUCTION

The main challenge in the analysis of multi-environment trials (MET) in plant breeding and variety testing is modelling and exploiting genotype-environment interaction. One of the most popular methods for this purpose is a regression of genotype performances in the individual environments on the environmental mean. This method was originally proposed by Yates and Cochran (1938) and later popularized by the seminal paper by Finlay and Wilkinson (1963). We will henceforth refer to this approach as Finlay-Wilkinson regression. In this regression, the environmental mean serves as an index for the environmental conditions. These conditions are determined by a large array of environmental variables, and it therefore seems natural to replace the environmental mean with measurable environmental covariates. Early treatments of such regression models for MET, also known as factorial regression (FR) (Denis, 1988), are found in Abou-El-Fittouh et al. (1969) and Freeman and Perkins (1971). The main challenge with FR is that the number of environmental covariates may be large, which makes the FR model very complex. This led to the idea to regress the environmental mean on covariates, instead of regressing the genotype-environment means on covariates separately for each genotype. This idea is so natural that it is hard to say who originally invented it. It is probably fair to say that the idea underlies and motivates the use of Finlay-Wilkinson regression, even in cases where observable environmental covariates are not used in the analysis (Piepho, 2022). A rigorous treatment of the idea in an MET context was first put forward by Hardwick and Wood (1972), who showed how to estimate the model parameters by the method of least squares. A partial least squares (PLS) approach to fit the same kind of model was proposed by Aastveit and Martens (1986), and this may be of particular interest when the number of covariates exceeds that of the environments. Van Eeuwijk (1992, 1995) pointed out that these models are related to what is known as redundancy analysis in psychology and as reduced rank regression in other applied areas such as engineering (Davies and Tso, 1982). An early mathematical treatment of the approach is found in Rao (1964).

All of the references considered so far assume that the genotype-environment classification is complete and that the residuals from the regression are independent with constant variance so that the ordinary least squares method provides optimal estimates. It must be acknowledged, however, that MET data are often unbalanced. Moreover, the variance-covariance structure needed to fully represent the experimental design, as well as to meet the objectives of the analysis, may deviate from the simple structure assumed for ordinary least squares, calling instead for a linear mixed model with additional random effects. This may, in fact, be one reason why these methods have not yet found very widespread use, despite an urgent need for such methods in an era where environmental information is becoming readily available and breeders are keenly interested in leveraging such information to make better selections and predictions (Xu, 2016; Cooper and Messina, 2021; Costa-Neto et al., 2021; Resende et al., 2021; Diepenbrock et al., 2022). The purpose of this paper, therefore, is to review the classical work based on ordinary least squares and to explore ways in which the approach can be extended to deal with unbalanced data and the need to use a mixed model framework.

The rest of the paper is organized as follows. In Section 2 we briefly recapitulate the Finlay-Wilkinson regression model and common methods for estimating it. Section 3 described the extension where the latent environmental score is regressed on environmental covariates and Section 4 considers the extension to more than one latent environmental score. In all three sections the residual is assumed to be independent with constant variance so that ordinary least squares can be applied. Our exposition in these sections mainly focuses on the models and only sketches methods of estimation. In Section 5 we consider the extension to mixed models and focus on those estimation approaches in the preceding sections which seem most suitable for this extension. For these select methods we then provide more details of the estimation steps. An example is presented in Section 6 to illustrate and compare the methods. The paper ends with a discussion in Section 7.

## 2 FINLAY-WILKINSON REGRESSION

The Finlay-Wilkinson model assumes that the response of different genotypes in varying environments can be modelled using the linear predictor

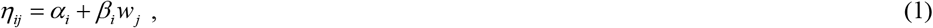

where *η*_*ij*_ is the expected performance of the *i*-th genotype (*i* = 1, …, *n*) in the *j*-th environment (*j* = 1, …, *m*), *α*_*i*_ and *β*_*i*_ are intercept and slope for the *i*-th genotype and *w* _*j*_ is a latent effect of the *j*-th environment. The slope *β*_*i*_ can be interpreted as a measure of sensitivity with small absolute values indicating stable responses over changing environments (Becker and Leon, 1988). The observed data are assumed to be genotype-environment means *y*_*ij*_, for which the model is *y*_*ij*_ = *η*_*ij*_ + *e*_*ij*_, where *e*_*ij*_ are independently and identically distributed residuals. For balanced data, Finlay and Wilkinson (1963) estimated *w*_*j*_ by the arithmetic mean of all observed genotype mean yields, *y*_*ij*_, i.e., they used the estimator 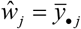. This, however, does not yield the least-squares fit of (1). The least squares fit can be obtained by a singular value decomposition (SVD) of the matrix 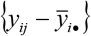, extracting the first singular vector for environments (Williams, 1952; Hardwick and Wood, 1972; Yan and Kang, 2003). Again, this assumes balanced data.

For unbalanced data, Digby (1979) proposed obtaining the least squares fit by alternating least squares, also known as criss-cross regression (CCR) (Gabriel and Zamir (1979), and Ng and Grunwald (1997; also see Ng and Williams, 2001) showed how to do this using nonlinear least squares. The model is not linear in the parameters, and some restriction on the parameters is needed for the multiplicative term *β*_*i*_*w*_*j*_. A further method initially replaces the empty cells of the two-way classification with initial values, then applies SVD to the completed table, re-estimates the empty cells from the fitted model, and so forth until convergence (Gauch and Zobel, 1990). We here focus on the other two methods because they are easily generalized to mixed models and a regression on covariates.

## 3 MODELLING THE ENVIRONMENTAL MEAN USING ENVIRONMENTAL COVARIATES

A downside of model (1) is that it cannot be used to predict the performance in unseen environments. If *w*_*j*_ can be replaced by an observable covariate, such predictions become possible. However, a single covariate rarely provides good predictions. Thus, a natural extension is to do a multiple regression on several covariates (Hardwick and Wood, 1972; Denis, 1988). Such an FR model quickly becomes very complex, because each treatment needs to have a separate regression coefficient for each environmental covariate.

For these reasons, it is desirable to consider more parsimonious alternatives. Specifically, one may consider regressing *w* _*j*_ on *p* observable covariates *x*_*jk*_ (*k* = 1, …, *p*), i.e.,

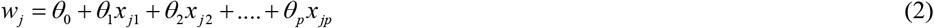

(Li et al., 2018; Guo et al., 2021). Importantly, this is just one multiple regression, instead of *n* multiple regressions for *n* genotypes. Inserting this into (1), we find

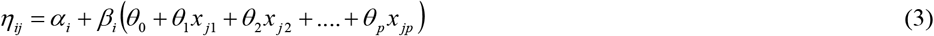

It is seen that the regression model is not linear in the parameters either, and there is an over-parameterization that needs to be resolved. Specifically, the intercept term *α*_*i*_ is fully confounded with the multiplicative term *β*_*i*_*θ*_0_. In fact, because of this confounding, we may drop the intercept term *θ*_0_ without loss of generality, as this term will then be absorbed by the intercept *α*_*i*_. The only caveat when dropping *θ*_0_ is that the model no longer involves a regression on the environmental mean. Instead, we have a regression on a synthetic covariate formed as a linear combination of observable environmental covariates. As this view is the most useful one for several of the methods considered below, especially when extending the models to comprise more than one synthetic environmental covariate (Section 4), we here also state the corresponding reparameterized (and equivalent) model explicitly. Thus,

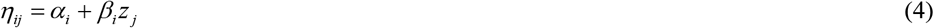

where

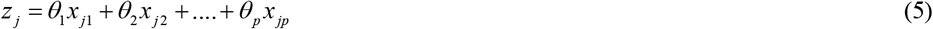

is a synthetic covariate. Several methods are possible to fit this regression model, and they differ in the way they deal with the over-parameterization. Here, we will consider several options, starting from simple but approximate methods assuming balanced data, and ending with an approach that will yield the least squares fit, also works for unbalanced data, and is readily extended to mixed models. Intermediate approaches give up optimality of the final estimate, with the benefit of a simplification of the pivotal step to find the values of the coefficients *θ*_*k*_ (*k* = 1,…, *p*) for the synthetic covariate *z* _*j*_. In this section, it will be assumed that all effects except the residual error term are fixed.

### 3.1 Balanced data

i. Consider the environmental averages based on (3):

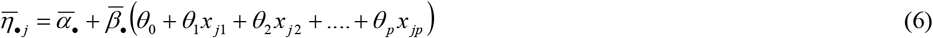

This model for environmental means suggests that a multiple regression of observed environmental means 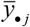 on the covariates provides estimates of slopes 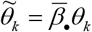 for covariates *x* _*jk*_ (*k* = 1,…, *p*) and the intercept 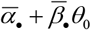. Without loss of generality, we may then use

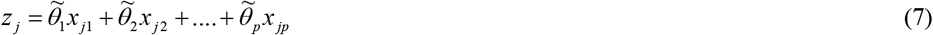

as our predictor for the environmental index in (4). This approach does not yield a least squares fit.
ii. Instead of using environmental means, we can obtain estimates *ŵ*_*j*_ of the latent environmental scores *w*_*j*_ in (1) using a least squares approach based on an SVD (Hardwick and Wood, 1972) and then use these estimates in place of *w*_*j*_ in (2) to estimate the regression parameters *θ*_1_,…,*θ* _*p*_, pretending subsequently that these estimates are known constants. This method does not provide a least squares fit.
iii. We may fit the FR model

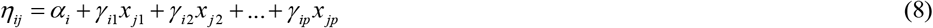

and subsequently subject the matrix of fitted terms 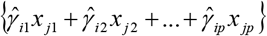 to an SVD. The first term of this decomposition provides the least squares fit for *β*_*i*_ *z* _*j*_ in (1) with *z*_*j*_ as given in (5) (Hardwick and Wood, 1972; Wood, 1976; Davies and Tso, 1982; van Eeuwijk et al., 1992). The least squares fit for *α*_*i*_ in (1) is that obtained from the fit of (8). This approach is also known as reduced rank regression or redundancy analysis.
iv. We fit (4) using the first factor extracted by partial least squares (PLS; Aastveit and Martens, 1986; Vargas et al., 1998), regarding the genotypic responses in an environment as a single multivariate response. PLS is usually performed scaling both the response variables and the covariates to zero mean and unit variance. For MET data, it is preferable to preserve the original scale of the genotypic responses. When using a multivariate PLS routine that allows scaling to be suppressed, this analysis may be obtained by scaling the covariates but not the responses before submitting the data to the PLS routine with the scaling option switched off.
v. We fit (4) directly by nonlinear least squares (Ng and Grunwald, 1997; Ng and Williams, 2001).

### 3.2 Unbalanced data

Among the five methods for balanced data reviewed in Section 3.1, method (i) cannot be used with unbalanced data. Method (v) works equally with unbalanced data. Methods (ii) to (iv) can be used with modification as described below.

(ii) Estimate *w*_*j*_ based on (1) using CCR (Dibgy, 1979; Gabriel and Zamir, 1979; Hadasch et al., 2018), then regress this estimate on *x* _*j*1_, *x* _*j* 2_, …, *x* _*jp*_ ; next replace *w*_*j*_ in (3) by its prediction based on this regression and fit the model parameters *α*_*i*_ and *β*_*i*_ by linear regression.
(iii) Fit the FR model (8) and obtain the matrix of fitted terms 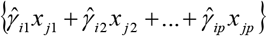 of all cells, including the ones with no data. Then proceed as in (iii) in Section 3.1.
(iv) We can use a method akin to the expectation-maximization (EM) method for fitting the AMMI model (Gauch and Zobel, 1990). The method starts by filling the empty cells of the genotype-environment classification with some plausible values, for example the genotype means. Then multivariate PLS is applied to the completed data and predictions are obtained to update the imputed values for the cells with missing data. This is repeated until predictions for the cells with missing data converge. For details, see Nelson et al. (1996). This EM method is implemented in the PLS procedure of SAS. It does not seem to be available in R, but is easily programmed using any PLS package for complete data.

## 4 MORE THAN ONE SYNTHETIC ENVIRONMENTAL COVARIATE

The model (4) may be extended as

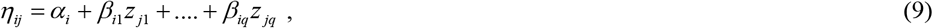

where *z* _*j*1_,., *z* _*jq*_ are the *q* synthetic environmental covariates and *β*_*i*1_,., *β*_*iq*_ are the corresponding genotype-specific slopes. This model may be estimated using the same methods as those in Section 3, excluding method (i). The only additional requirement is that estimability constraints, such as orthogonality, need to be imposed on estimates of both *z* _*j*1_,., *z* _*jq*_ and *β*_*i*1_,., *β*_*iq*_, when fitting explicit regressions on observable environmental covariates of the form

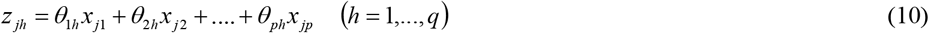

Note that we have dropped the intercept terms *θ*_0*h*_, as these will be confounded with the genotype-specific intercepts. Where an SVD is used (methods iii and iv), constraints are automatically imposed as part of the decomposition, and we here extract the first *q* multiplicative terms. Where CCR or nonlinear least squares are used (methods ii and v), the constraints need to be actively imposed on the *q* multiplicative terms, e.g., by subjecting current estimates to an SVD on each iteration (Hadasch et al., 2018).

## 5 RANDOM-EFFECTS EXTENSIONS OF THE MODEL

If the regression model comprises random effects, it is referred to as a mixed model. There are various reasons why random effects may be needed with MET. Here, we will generally regard environments as a random factor, assuming that trial environments represent a random sample from a target population of environments (TPE). The assumption or random environments is at the heart of different concepts of phenotypic stability, among which Finlay-Wilkinson regression and its extensions are one of the most prominent examples (Becker and Leon, 1988; Piepho, 1998). Furthermore, this assumption is needed to project the model to unseen environments in the TPE (Buntaran et al., 2021). This assumption will therefore be made throughout this section. A second reason for introducing random effects is to allow genetic markers to be used for modelling genotypic effects as in genomic prediction (Meuwissen et al., 2001; Bernardo and Yu, 2007). In this case, which is considered in Section 5.2, the genotype factor needs to be modelled as random as well.

### 5.1 All parameters in *η*_*ij*_ are fixed

When modelling the variance over random environments, there are different aspects calling for random-effects modelling. For example, observed data may display heterogeneity of variance between genotypes in the deviations from the regression line. This treatment-specific variance has been proposed by Eberhart and Russell (1966) as an additional stability parameter to the regression coefficient *β*_*ik*_. Furthermore, a covariance must usually expected between genotypes in the same environment due to a residual main effect for the shared environment. This can be modelled, e.g., by a random environmental main effect. Thus, our model for the mean response *y*_*ij*_ of the *i*-th genotype in the *j*-th environment may be written as

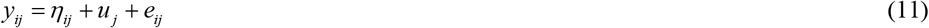

where *η*_*ij*_ is as defined in (9), *u*_*j*_ is the random year main effect with variance 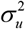, and *e*_*ij*_ is a random with stability variance 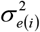 for the *i*-th genotype (Eberhart and Russell, 1966; Shukla, 1972).

Of course this is just one possible mixed-model extension. The random deviations from the regression in (9) may be modelled in different ways depending on the data structure. For example, so far, we have assumed that the model is fitted to genotype-environment mean responses *y*_*ij*_. Alternatively, replicate plot data may need to be modelled, requiring additional random design effects. With perennial crops or in long-term trials, it may be necessary to model serial correlation of observations on the same plot (Macholdt et al., 2022). All of these mixed-model extensions are straightforward with the general approaches suggested in this section. The distinction between balanced and unbalanced data becomes a moot point here, because the variance-covariance structure used in the mixed model usually implies that ordinary least squares estimation and the use of simple arithmetic means for genotypes or environments in the estimation process are not usually a good option even when the data is balanced. Hence, we focus on the methods presented in Section 3.2. We do not present an adaptation of the PLS method (iv) to mixed models, because we are not aware of any proposed method or package that would provide this. Thus, we focus on methods (ii), (iii) and (v).

(ii) Following Nabugoomu et al. (1999), the CCR approach of Digby (1979) is easily extended in a mixed model framework. In the criss-step, estimating *θ*_*hk*_ (*k* = 1,…, *p*;*h* = 1,…, *q*), we fix the variance parameters at their current estimates because this usually has fewer parameters to be estimated as fixed effects than the cross-step. These variance parameters are re-estimated in the cross-step, estimating *α*_*i*_ and *β*_*ih*_ (*i* = 1,…, *n*;*h* = 1,…, *q*), using residual maximum likelihood. For an application of this method, see Macholdt et al. (2022).
(iii) We may fit FR model (8) under our assumed random-effects specification in (11) using residual maximum likelihood. From the fitted model we obtain the matrix of fitted terms 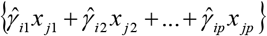 of all cells, including ones with missing data. This matrix is subjected to an SVD to estimate the parameters in (9) and (10). Net, we compute *z*_*jk*_ according to (10) refit (9) to estimate *α*_*i*_ and *β*_*ih*_ (*i* = 1,…, *n*; *h* = 1,…, *q*). As a refinement we can consider a weighted SVD using the approach of Hadasch et al. (2018), taking into account the variance-covariance matrix of predictions 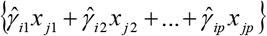.
(v) Fit (9) with (10) directly using full maximum likelihood (ML) [e.g., using NLMIXED in SAS; also see Piepho (1999)]. Note that we can not use residual maximum likelihood (REML) because the model is non-linear in the parameters for *η*_*ij*_ and hence the fixed effects cannot be removed by linear contrasts as in REML. Full ML does not account for the degrees of freedom and hence leads to more biased variance parameter estimates than REML with linear mixed models. These problems are expected to carry over to the nonlinear mixed model (9) with (10). There is no REML equivalent because the model is intrinsically nonlinear. The important consequence is that in order to avoid the bias issues with full ML, we need to resort to approximate methods such as (ii) to (iv) that make use of REML.

Apart from the above modifications for mixed models, there is always the option to apply methods (ii) to (v) in the same way as in Sections 3.2 and 4, but applying these to fitted means *η*_*ij*_ using a suitable mixed model for the variation of the observed data around these means. The main objective of applying those methods is to get coefficients *θ*_*kh*_ so we can compute the synthetic covariates *z* _*jh*_ in (10). Once these are available, we simply treat them as if they were known covariates. While this only constitutes an approximation and can not be optimal, partly because the fact is ignored that *z* _*jh*_ involves coefficients estimate from the data, it does have the advantage of simplicity. So despite imperfections, this may be the most easily implemented approach in practice.

### 5.2 Some or all genotypic parameters in *η*_*ij*_ are random

When the number of genotypes is large, it may be advantageous to fit both *α*_*i*_ and *β*_*ih*_ as random. This may be particularly worthwhile when marker information is available so kinship information can be used to perform genomic prediction by GBLUP (Bernardo and Yu, 2007). Furthermore, genotypes need to be modelled as random in case TPE is stratified into zones and we want to borrow strength across zones (Buntaran et al., 2021).

It is not as straightforward as it may seem to simply switch from fixed to random genotypes. This is because mixed model packages assume that all random effects have an expected value of zero. This is not a realistic assumption if we postulate the Finlay-Wilkinson model (1) and its extensions as considered so far, such as model (4). To see this, assume the response indeed obeys (4), i.e. we only have one synthetic environmental covariate, and we now take both the intercept *α*_*i*_ and the slope *β*_*i*_ to be random. We can assume that *α*_*i*_ has expected value *μ*_*α*_ and hence set *α*_*i*_ = *μ*_*α*_ + *a*_*i*_ with *E*(*a*_*i*_) = 0 and 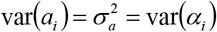. This can be fit with a linear mixed model package because the intercept enters the model linearly. We may consider the same approach for the slope, setting *β*_*i*_ = *μ*_*β*_ + *b*_*i*_ with *E*(*b*_*i*_) = 0 and 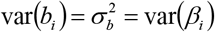. The model (4) may then be re-written as

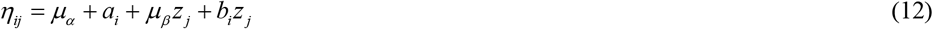

The main challenge with this model is that the latent score, *z*_*j*_, now appears both in the fixed-effects term *μ*_*β*_ *z* _*j*_ and the random effects term *b*_*i*_ *z* _*j*_ (Piepho, 1999). The key point is that we cannot assume *μ*_*β*_ = 0 without loss of generality, because there is no fixed environmental main effect in (12) that would absorb this term. Also, *μ*_*β*_ *z* _*j*_ is a multiplicative regression term that may be worth fitting explicitly rather than absorbing into an environmental main effect. Furthermore, to ensure invariance to shift-scale transformations of the observable covariates, we need to allow for a covariance between intercept and slope, i.e., cov(*a*_*i*_, *b*_*i*_) = σ _*ab*_ (Piepho, 1999). Either genotypes are modelled as independent, or they are modelled as correlated by kinship for GBLUP (Resende et al., 2021).

Two methods seem feasible for fitting this model and its extensions to more than one synthetic environmental covariate. We may apply a stage-wise approach by which we first model all parameters in (9) and (10) as fixed, using either of the methods (ii) to (iv). This provides estimates of *w*_*j*_, which may then be held fixed and used in place of *w*_*j*_ in (12), using REML to fit 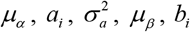, and 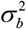. Extension to models with more than one latent environmental scale is straightforward. The other method is alternating weighted least squares, adapting the approach of Nabugoomu et al. (1999).

A further option is to model the genotype-specific intercepts *a*_*i*_ in (12) as fixed, while modelling the slopes *b*_*i*_ as random, thus obviating the need to fit a covariance between intercept and slope (Piepho and Ogutu, 2002). This approach will be the focus of the remainder of this section. We further add a fixed environmental main effect *ε*_*j*_ that absorbs *μ*_*β*_ *w*_*j*_. Hence, the only random effect in *η*_*ij*_ is the slope *b*_*i*_, and the model can be written as

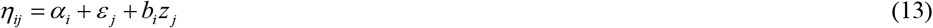

with *E*(*b*_*i*_) = 0 and 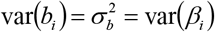. This model is readily extended to comprise several synthetic environmental covariates:

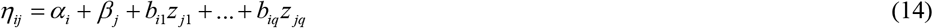

It will now be shown how (14) can be cast as a random-coefficient FR model for the observed covariates, in which a factor-analytic (FA) variance-covariance structure is assumed for the random regression coefficients. Let ***x*** = (*x*_*j*1_, *x*_*j*2_,…, *x*_*jp*_)^*T*^, so that 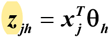 with ***θ***_*h*_ = (*θ*_1*h*_, *θ*_2*h*_, …,*θ*_*ph*_)^*T*^. Moreover, re-write the random terms in (14) as

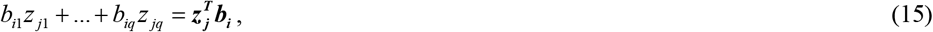

where ***z*** = (*z*_*j*1_, *z*_*j*2_,…, *z*_*jq*_)^*T*^ and ***b***_***i***_ = (*b*_*i* 1_, *b*_*i* 2_,…,*b*_*iq*_)^*T*^. Now first consider a random coefficient regression of the form

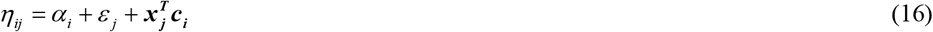

where ***c***_***i***_ = (*c*_*i*1_, *c*_*i*2_, …, *c*_*ip*_)^*T*^ with *E*(***c***_***i***_) = **0** and var(***c***_***i***_) = **∑**_***c***_ (Longford, 1995). Next assume that we approximate the variance-covariance matrix **∑**_***c***_ by the model

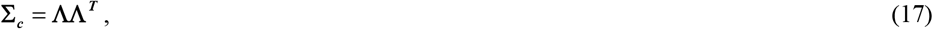

where **Λ** = {*λ*_*kh*_} is a *p* × *q* matrix of factor loadings (Buntaran et al., 2021; Tolhurst et al., 2022). This amounts to an FA structure of order *q* for the regression coefficients ***c***_***i***_ without residual effects and associated specific variances. To ensure estimability, we impose the constraints *λ*_*kh*_ = 0 for *h* > *k* (Jennrich and Schluchter, 1986). This structure is straightforward to fit using some mixed model packages. For example, in SAS this is the FA0(*q*) structure. With this approximation, the regression term can be re-written as

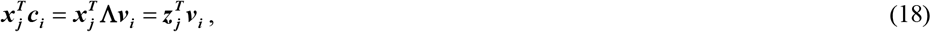

where ***v***_***i***_ = (*v*_*i*1_, *v*_*i*2_, …, *v*_*iq*_)^*T*^ with *E*(***v***_***i***_) = **0** and var(***v***_***i***_) = ***I***_***q***_, and

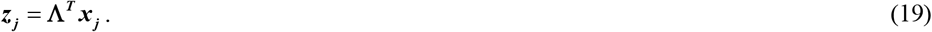

Comparing coefficients between (15) and (18), it emerges that, apart from a difference in scaling, we can equate ***v***_***i***_ with ***b***_***i***_ and **Λ** with **Θ** = {*θ*_*kh*_}. The important practical consequence of this observation is that we can simply fit (16) with an FA0(*q*) structure for ***c***_***i***_, extract the estimate of **Λ** and use this in (19) to compute the *q* synthetic covariates ***z***_***j***_ for any environment *j*, including unobserved ones, so long as we have their covariate values ***x***_***j***_. Furthermore, with this approach we can subsequently set var(***b***_***i***_) = ***I***_***q***_, if genotypes are to be modelled as random. The full model (12) can then be re-fitted, taking intercepts *a*_*i*_ as random as well and allowing for a covariance among *a*_*i*_ and ***b***_***i***_. Alternatively, we may model genotypes as fixed for the final analysis and regard the random-effects analysis as a convenient intermediate tool for obtaining ***z***_***j***_. In either case, the fixed regression term 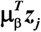 can now be estimated explicitly in the final step, rather than absorbing this into an environmental main effect as in (13). This approach for the final step is approximate, as it treats ***z***_***j***_ as if these were observable covariates not involving parameters.

## 6 EXAMPLE

To illustrate the random-effects modelling discussed in Section 5, we consider the lettuce data reported in van Eeuwijk (1992) as means per genotype and environment. The response is nitrate concentration. The data is balanced and stems from a single replicated trial, in which eight genotypes were evaluated at 18 points in time, which are regarded as environments for our analyses. The data comprise eight observed environmental covariates, which are scaled here to zero mean and unit variance for all regression analyses. We start by fitting model (16), dropping the covariate terms, leaving the structure *α*_*i*_ + *β*_*j*_. Our baseline model in Table 1 has independent (ID) residual effects *e*_*ij*_ with constant variance. Replacing this by an AR(1) model for serial correlation of observations on the same genotype across the 18 times (environments) leads to a substantial drop in the Akaike Information Criterion (AIC) (Wolfinger, 1996), indicating that serial correlation is important for this repeated measures data. Next, we add random effects 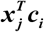 as in model (16) and fit an FA0(1) structure for the covariance among random slopes ***c***_***i***_. This leads to a further marked drop in AIC, showing that the covariates are important. The variance parameter estimates of the FA0(1) model are given in Table 2 for illustration. The serial correlation of 0.40 is non-negligible. The estimated loadings are used to compute the synthetic variable *z*_1_ = *λ*_1_*x*_1_ + … + *λ*_8_*x*_8_ (see eq. 19). With this, we then fit model (4), adding a random main effect 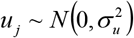 for environments and using the AR(1) model for the residual. Regression with this covariate has a significant interaction with genotype (Table 3), indicating there are differences in sensitivity. The sensitivities (slopes) as well as the intercepts for the eight genotypes are shown in Table 4. The variance explained is substantial, as a comparison of the models with and without covariate shows (Table 5). Adding a second latent factor using FA0(2) leads to a further improvement in fit (Table 1). Detailed results are omitted here for brevity.

**Table 1:**
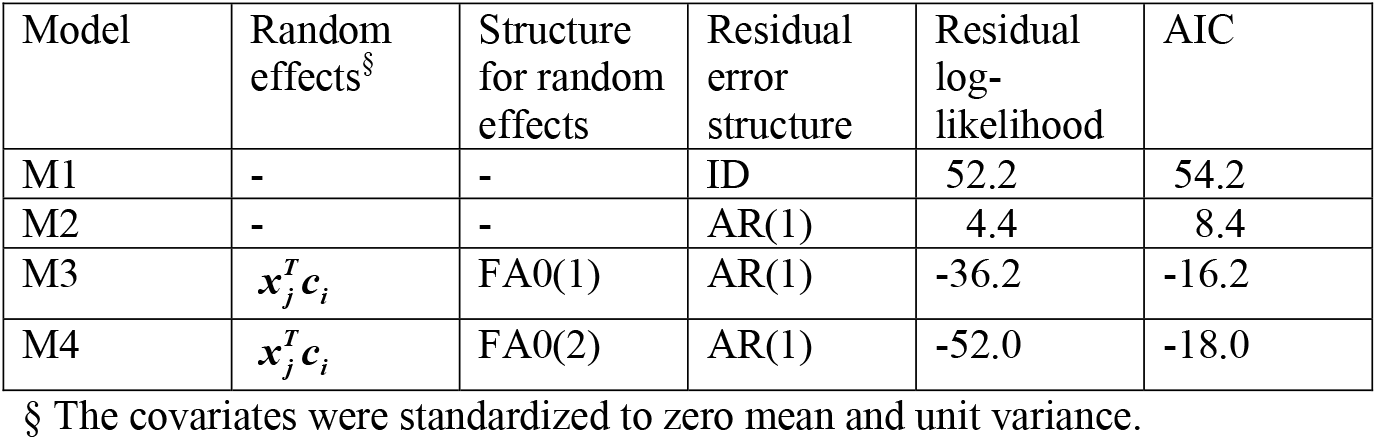
Model selection of covariance structure for lettuce data using fixed effects *α*_*i*_ + *ε*_*j*_.

| Model | Random effects<sup>§</sup> | Structure for random effects | Residual error structure | Residual log-likelihood | AIC |
| --- | --- | --- | --- | --- | --- |
| M1 | - | - | ID | 52.2 | 54.2 |
| M2 | - | - | AR(1) | 4.4 | 8.4 |
| M3 | $\mathbf{x}_j^T \mathbf{c}_i$ | FA0(1) | AR(1) | -36.2 | -16.2 |
| M4 | $\mathbf{x}_j^T \mathbf{c}_i$ | FA0(2) | AR(1) | -52.0 | -18.0 |

§ The covariates were standardized to zero mean and unit variance.

**Table 2:** REML estimate of variance parameters in Model M3 of Table 1 (*q* = 1) for lettuce data.

| Parameter | Estimate | S.E. |
| --- | --- | --- |
| $\lambda_1$ | 0.009574 | 0.02459 |
| $\lambda_2$ | -0.06102 | 0.06146 |
| $\lambda_3$ | 0.2711 | 0.1355 |
| $\lambda_4$ | -0.2763 | 0.1173 |
| $\lambda_5$ | 0.07636 | 0.07623 |
| $\lambda_6$ | 0.01464 | 0.03586 |
| $\lambda_7$ | 0.09994 | 0.06392 |
| $\lambda_8$ | 0.07464 | 0.04965 |
| $\rho$ | 0.3999 | 0.1128 |
| $\sigma^2$ | 0.02897 | 0.005395 |

**Table 3:** Wald-type F-tests for regression with synthetic variable *z* obtained from FA0(1) model. Lettuce data.

| Effect | Numerator d.f. | Denominator d.f. | F-value | p-value |
| --- | --- | --- | --- | --- |
| Genotype | 7 | 28.1 | 74.25 | <.0001 |
| $z$ | 1 | 18.5 | 6.58 | 0.0192 |
| Genotype $\times z$ | 7 | 79.9 | 6.68 | <.0001 |

**Table 4:** Intercepts and slopes of regression with a single synthetic environmental covariate *z*_1_ = *λ*_1_*x*_1_ + … + *λ*_8_*x*_8_ computed using different methods (FA, PLS, CCS, SVD). Lettuce data.

| Genotype | FA |  | PLS |  | CCR |  | SVD |  |
| --- | --- | --- | --- | --- | --- | --- | --- | --- |
|  | Intercept | Slope | Intercept | Slope | Intercept | Slope | Intercept | Slope |
| DM | 3.0830 | 4.2928 | 3.0875 | 0.2329 | 0.2034 | 1.9167 | 1.8737 | 5.7769 |
| GT | 2.4156 | 2.0209 | 2.4188 | 0.1054 | 1.3101 | 0.7356 | 1.7816 | 3.0054 |
| Ls | 2.7136 | 1.1955 | 2.7115 | 0.08166 | 2.0877 | 0.4153 | 2.1631 | 2.5973 |
| Pa | 3.4633 | 3.9315 | 3.4601 | 0.1842 | 0.9184 | 1.6922 | 2.4296 | 4.9122 |
| Pi | 2.8687 | 3.2520 | 2.8673 | 0.1589 | 0.9482 | 1.2767 | 2.0449 | 3.9028 |
| RW | 2.4862 | 0.7351 | 2.4832 | 0.02304 | 2.3516 | 0.08801 | 2.2234 | 1.2035 |
| Tr | 1.7665 | 2.4550 | 1.7700 | 0.1555 | -0.1831 | 1.2986 | 0.7840 | 4.6952 |
| Wi | 2.6674 | 1.4932 | 2.6688 | 0.07303 | 1.8017 | 0.5760 | 2.1178 | 2.5985 |
| Standard error | 0.1209 | 1.1531 | 0.1024 | 0.04263 | 0.5334 | 0.3485 | 0.1914 | 0.8099 |

**Table 5:** Variance parameter estimates (REML) for intercept only model and for regression with a single synthetic environmental covariate *z*_1_ = *λ*_1_*x*_1_ + … + *λ*_8_*x*_8_ computed using different methods (FA, PLS, CCS, SVD), as well as FR using the observed covariates *x*_1_-*x*_8_. Lettuce data.

| Parameter | Description | Without covariates | | With covariate $z_1$ | | | | | | | | With covariates $x_1$-$x_8$ | |
| --- | --- | --- | --- | --- | --- | --- | --- | --- | --- | --- | --- | --- | --- |
|  |  |  |  | FA |  | PLS |  | CCR |  | SVD |  | FR |  |
|  |  | Estimate | S.E. | Estimate | S.E. | Estimate | S.E. | Estimate | S.E. | Estimate | S.E. | Estimate | S.E. |
| $\sigma_u^2$ | Variance of environmental main effect | 0.1959 | 0.07176 | 0.1466 | 0.05305 | 0.1277 | 0.04645 | 0.1311 | 0.04752 | 0.06737 | 0.02510 | 0.07649 | 0.03738 |
| $\rho$ | Autocorrelation [AR(1) model] | 0.6190 | 0.08591 | 0.4131 | 0.1061 | 0.3328 | 0.1070 | 0.4125 | 0.1072 | 0.4014 | 0.1099 | 0.5421 | 0.2480 |
| $\sigma^2$ | Residual error variance | 0.06686 | 0.01529 | 0.02943 | 0.005464 | 0.03227 | 0.005426 | 0.02965 | 0.005544 | 0.03666 | 0.006855 | 0.03294 | 0.01568 |

For comparison, we also used CCR, PLS and SVD to obtain the synthetic environmental covariate *z*_1_ (Tables 4 and 5). The results are broadly similar in terms of the ranking of sensitivities (Table 4), even though there are differences of scale. The variance explained is also comparable between the different approaches (Table 5).

Table 6 gives an overview of the fits obtained by the different methods and models when using full maximum likelihood (ML), thus permitting comparison of models with different fixed effects (Wolfinger, 1996). This also comprises model estimates based on SVD using the fitted factorial regression model (option iii). The AIC values in Table 6 reveal that CCR provides the best fit among the methods using a single synthetic covariate *z*_1_, closely followed by FA. When using two synthetic covariates (*z*_1_, *z*_2_), the best fit is obtained with the FA approach. That fit is also better than FR, which has a large number of parameters. We have not implemented CCR with two covariates, but this is expected to give comparable fit.

**Table 6:**
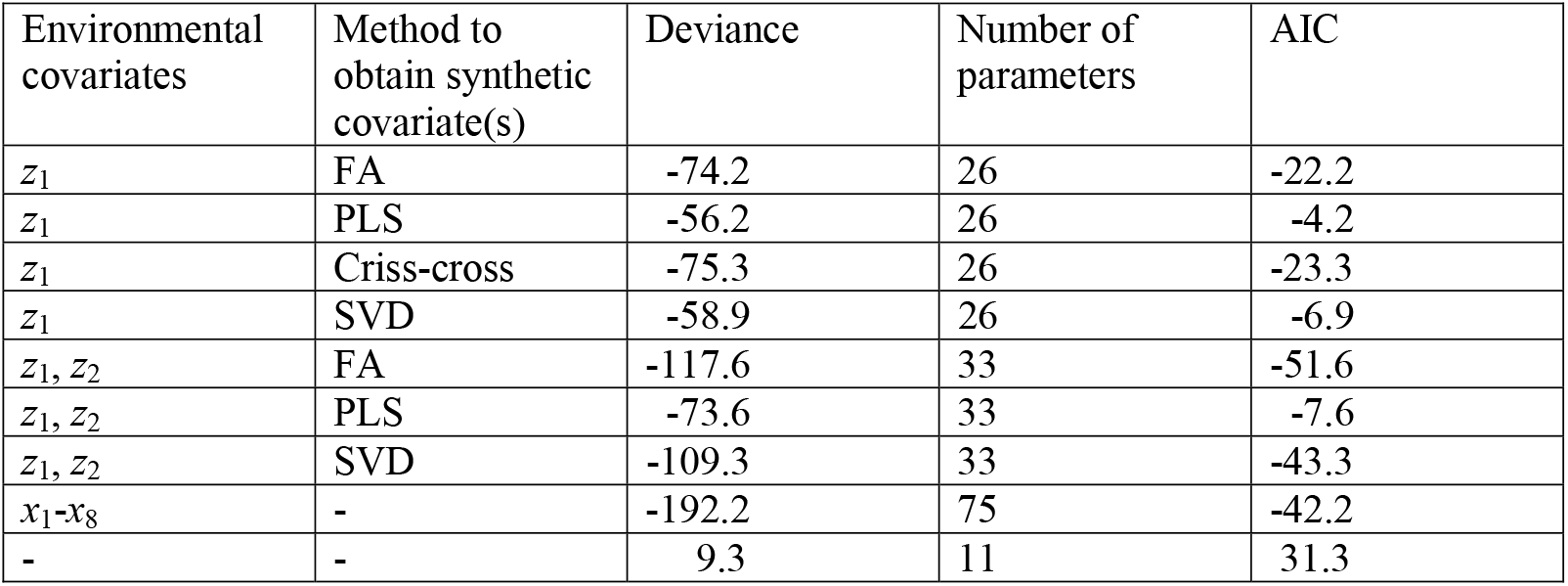
Deviance (full likelihood) and AIC for different models. Lettuce data.

## 7 CONCLUDING REMARKS

Among the different methods for obtaining synthetic environmental covariates to be used in model (4), the FA approach that models genotype-specific regression coefficients ***c***_***i***_ for the observed environmental covariates ***x***_***j***_ as random did best in the example. Moreover, the FA approach is much easier to implement than the closest competitor, CCR, especially when use of more than one synthetic covariate is considered. Both if these methods for obtaining the synthetic covariates use a mixed model that is commensurate with the model finally fitted once the synthetic covariates are obtained. This explains why they perform best. By contrast, The PLS and SVD approaches do not fully take the final mixed model into account when estimating the synthetic covariates, as they essentially assume i.i.d. residual errors. The main advantage of the PLS approach over all other approaches considered here is that it can handle a larger number of observed covariates.

Using synthetic environmental covariates *z*_*jh*_ allows fitting more parsimonious regression models than when regressing directly on observed environmental covariates. Not only does this allow a more efficient analysis, but it also facilitates interpretation. When there are *p* observed environmental covariates *x*_*jk*_, FR involves *p* regression coefficients for each genotypes, which may be difficult to interpret when *p* is large. By contrast, if these *p* observed covariates are used to form a single synthetic covariate, a single regression coefficient is involved per genotype, and this can be interpreted as a sensitivity parameter, as with Finlay-Wilkinson regression.

Our FA model in Section 5.2 has similarities with the model proposed by Tolhurst et al. (2022). Those authors initially model environments as fixed [see their model (1)], whereas we model environments as random throughout. When extending their model to allow for environmental covariates, the authors do consider a random environmental effect for deviations from the fixed-effects regression on covariates ***x***_***j***_ to model the environmental main effect. The regression on observed covariates ***x***_***j***_ would absorb our term *μ*_*β*_ *z*_*j*_. However, their model does not employ the more parsimonious regression on the synthetic covariates in the fixed part of the model, which is a major difference from our model. Also, synthetic variables ***z***_***j***_ do not feature explicitly in the random part of their model, but they do so implicitly. Another difference is that Tolhurst et al. (2022) model genotypes as random throughout. Specifically, they fit random intercepts *a*_*i*_ (our notation) throughout, allowing for a covariance with slopes ***c***_***i***_. By contrast, in our example we only fit random effects for slopes ***c***_***i***_, while modelling intercepts *a*_*i*_ as fixed, thus obviating the need to fit the covariances among *a*_*i*_ and ***c***_***i***_, which can be numerically challenging (Buntaran et al., 2021). The primary purpose of fitting our random-effects regression on observed environmental covariates is to estimate coefficients for the synthetic environmental covariates ***z***_***j***_ in eq. (19). This is just an intermediate step before fitting the final model, which may have fixed genotypic slopes for the regression on ***z***_***j***_.

## CONFLICT OF INTEREST

The authors declared no potential conflicts of interest with respect to the research, authorship and/or publication of this article.

## AUTHOR CONRIBUTIONS

All work was conducted by the author.

## DATA AVAILABILITY STATEMENT

The data used in this paper is available in literature and is included with the full computer code that allows reproducing all results.

## ACKNOWLEDGEMENTS

The author thanks the German Research Foundation (DFG) for financial support (grant PI 377/20-2).

## SUPPORTING INFORMATION

Additional supporting information can be found online in the Support-ing Information section at the end of this article.

## Notes

**Funding statement** This research was supported by DFG grant PI 377/20-2

**Conflict of interest** The author declares no conflict of interest

### Competing Interest Statement

The authors have declared no competing interest.

